# Late onset of striatal projection neuron hyperexcitability in *Fmr1*-/y mice

**DOI:** 10.1101/2025.06.21.660889

**Authors:** Lars Nelson, Michael Janeček, Michael Matarazzo, Yi-Chun Shih, Rui T. Peixoto

## Abstract

Fragile X Syndrome (FXS), the most common genetic cause of intellectual disability and autism spectrum disorder (ASD), results from silencing of the *FMR1* gene and consequent loss of Fragile X Messenger Ribonucleoprotein (FMRP). FMRP deficiency disrupts neural development, leading to behavioral and motor deficits associated with striatal dysfunction. While structural and functional abnormalities in striatal projection neurons (SPNs) have been observed in adult *Fmr1* knockout (KO) mice, their developmental onset and contribution to early FXS pathophysiology remain unknown. In this study, we examined the postnatal maturation of SPN in the dorsomedial striatum (DMS) of *Fmr1* KO mice, assessing glutamatergic synaptic inputs and intrinsic excitability. During postnatal development, *Fmr1* deficient SPNs in DMS display normal synaptic and intrinsic properties, consistent with typical maturation. In contrast, by P60, SPNs of mice exhibit pronounced hyperexcitability, characterized by increased membrane resistance, reduced rheobase, and slower action potential kinetics. These perturbations affect both Dopamine 1 receptor-expressing (D1-SPN) and D2 receptor-expressing (D2-SPN) SPNs, though some action potential dynamics are selectively impaired in D1-SPNs. Chronic aripiprazole treatment, a widely prescribed therapy for FXS-related symptoms, fails to normalize SPN excitability, highlighting its limited efficacy in addressing core SPN dysfunction. Our findings reveal a late-onset hyperexcitability in DMS SPNs of *Fmr1* KO mice, suggesting a progressive emergence of striatal neuron abnormalities over development. These results underscore the importance of developmental timing in FXS pathophysiology and emphasize the need for targeted interventions to address striatal circuit dysfunction.

## Introduction

Fragile X Syndrome (FXS), the most common inherited cause of intellectual disability and autism spectrum disorder (ASD), results from disruption of the 5′ untranslated region of the *FMR1* gene on the X chromosome ^1^. This disruption typically involves a CGG trinucleotide repeat expansion, leading to hypermethylation and transcriptional silencing of *FMR1* and a consequent loss of Fragile X Messenger Ribonucleoprotein (FMRP). FMRP, an RNA-binding protein abundantly expressed in the central nervous system, regulates the translation of numerous mRNA targets critical for synaptic plasticity and neural development ^2,3^. Its absence leads to a spectrum of behavioral and neurological impairments, including language delays, sensory hypersensitivity, irritability, hypotonia and a high prevalence of seizures, reflecting widespread neurological dysfunction ^4^. Mouse models of FXS have provided critical insights into the role of FMRP in cortical circuit development and function, with loss of *Fmr1* inducing abnormal neuronal excitability, synaptic plasticity, and long-range connectivity ^5–9^. Many of these deficits emerge early in development, with patterns of abnormal cortical activity already observed during the first postnatal weeks ^10–12^. Recent studies have also observed deficits in striatal circuits in adult *Fmr1* knockout (KO) mice, suggesting a potential role for the striatum in the motor and behavioral symptoms of FXS ^13–19^. However, whether striatal dysfunction is associated with the early pathogenesis of FXS remains unknown.

The striatum, the principal input structure of the basal ganglia, integrates diverse afferent inputs organized into distinct functional domains ^20^. The dorsal striatum is divided into the dorsomedial striatum (DMS), implicated in goal-directed behavior, and the dorsolateral striatum (DLS), associated with motor control. The ventral striatum, including the nucleus accumbens (NAc), mediates reward processing and emotional regulation. Deficits in these striatal functions align with core symptoms of autism, implicating striatal dysfunction in ASD ^21^, and neuroimaging studies consistently reveal hypertrophy and altered connectivity of striatal regions in individuals with ASD ^22–28^. Furthermore, striatal neurons exhibit one of the highest expression rates of ASD risk genes ^29,30^. Many of these genes, particularly those associated with synaptic maturation and function, are dynamically regulated in striatal neurons during the first postnatal weeks ^31^, further pointing to a convergence of ASD genetic risk in striatal circuit maturation. The activity of striatal circuits are primarily driven by glutamatergic input onto striatal projection neurons (SPNs), which express either dopamine D1 (D1-SPN) or D2 receptors (D2-SPN) ^20^. The functional balance between these two SPN populations is crucial for motor control and cognitive processes, with disruptions in striatal circuit activity linked to severe symptoms in a wide range of psychiatric and neurodevelopmental disorders, including ASD ^21,32–35^.

In adult *Fmr1*-/y mice, FMRP deficiency induces complex, region-specific alterations in striatal circuits. In the NAc, loss of FMRP impairs synaptic plasticity of glutamatergic synapses in SPNs ^16,18^. Additionally, NAc SPNs of *Fmr1*-/y mice exhibit altered intrinsic properties, with opposing changes in membrane excitability and action potential dynamics in D1-SPNs versus D2-SPNs^36^. These findings indicate a cell-type-specific role for FMRP in regulating properties of mature SPNs, with potential deleterious implications for striatal circuit function. In the DLS, the loss of FMRP alters dendritic structure and synaptic density ^13^. While an initial study reported subtle reductions in stubby dendritic spines across SPNs ^15^, subsequent analysis of separate SPN populations revealed a predominant increase in synaptic density in D1-SPNs, with no significant changes observed in D2-SPNs ^13^. In contrast, the effects of FMRP loss in the DMS remain poorly characterized, with the exception of one study that found no changes in dendritic spine density in DMS SPNs of adult *Fmr1*-/y mice ^15^. However, whether DMS D1-SPNs and D2- SPNs are differentially affected or exhibit functional abnormalities remains unknown. This gap in knowledge is particularly significant, as structural deficits in the head of the caudate—analogous to the DMS in rodents—have been reported in individuals with FXS ^37^ and are among the most recurrent findings in imaging studies of ASD ^23–28^. Additionally, FMRP expression peaks in the striatum during perinatal periods ^38^, suggesting a crucial role in the early maturation of SPNs. Cortical dysfunction in *Fmr1*-/y mice emerges during the first postnatal week ^10–12^, a period marked by tight developmental coupling between cortical and striatal circuits ^31,39,40^. This raises the possibility that SPN dysfunction arises early, driven by disruptions in both local and afferent developmental processes. However, the developmental trajectory of SPN deficits in *Fmr1*-/y remains unknown. Addressing whether SPN deficits arise during early postnatal development is critical for identifying effective windows of opportunity for therapeutic intervention.

Despite the high prevalence and significant disease burden of FXS, no prophylactic treatments have been developed ^41^. Aripiprazole, an atypical antipsychotic that is commonly prescribed to manage irritability and aggression in individuals with FXS ^42^. Given its pharmacological profile, aripiprazole can modulate presynaptic dopamine release by binding to the presynaptic D2 receptor as well as modulate D2-SPN activity. In addition, aripiprazole binds to the 5HT2A receptor, DRD3 and adrenergic receptor. Due to the broad pharmacological profile of aripiprazole, it is unclear whether the effects of aripiprazole are due to direct modulation of SPNs. However, its precise mechanisms of action and effectiveness in addressing the neural deficits caused by FMRP loss remain poorly understood. Notably, while aripiprazole can alleviate some behavioral symptoms, it is associated with significant adverse effects ^43,44^, highlighting the need for further investigation into its impact on striatal circuits and the development of alternative therapeutic strategies.

To address these gaps, we investigated the developmental trajectory of glutamatergic synaptic inputs and intrinsic excitability of DMS SPNs in *Fmr1*-/y mice across early postnatal (P14-15) and adult (P60) stages. In addition, we assessed the potential for chronic aripiprazole treatment to ameliorate intrinsic SPN dysfunction in adult *Fmr1*-/y mice. Our findings reveal a striking late-onset hyperexcitability of SPNs in the DMS that affects both D1-SPNs and D2-SPNs and is not rescued by chronic aripiprazole treatment. These results underscore the importance of developmental timing in FXS striatal pathophysiology and suggest that aripiprazole does not target core physiological abnormalities of SPNs caused by *Fmr1* deletion.

## Results

### Normal glutamatergic synaptic input in DMS D1- and D2-SPNs of *Fmr1*-/y mice at P15

To determine whether *Fmr1* deletion affects the early postnatal development of glutamatergic synapses onto DMS D1- and D2-SPNs, we recorded AMPAR-mediated miniature excitatory postsynaptic currents (mEPSCs) in acute brain slices from male *Fmr1+/y* or *Fmr1−/y* mouse pups carrying a *Drd1a-tdTomato* allele. Recordings were performed at P15, a developmental stage marked by rapid maturation of SPNs and analogous to early infancy in humans, when ASD symptoms typically begin to emerge ^39,45^. Expression of tdTomato was used for identification of D1-SPNs (Figure 1A). Importantly, the tdTomato-negative cell population includes striatal interneurons, but these represent a small fraction (<5%) of total striatal neurons ^46^ and we further excluded them from analysis based on their specific membrane resistance and capacitance properties (see Methods). For the remainder of this study, tdTomato-positive neurons are referred to as D1-SPNs, and tdTomato-negative neurons as D2-SPNs. To measure AMPAR miniature excitatory postsynaptic currents (mEPSC) we performed whole-cell recordings in the presence of the voltage-gated sodium channel blocker tetrodotoxin (TTX) with membrane potential clamped at -70mV. Quantification of mEPSC frequency revealed no significant difference between genotypes in either D1- or D2-SPNs (Figures 1C-D, Genotype: F_1,59_ = 0.0571, p = 0.812, η2 = 0.000946, 95% CI[-0.202, 0.257]; SPN subtype: F_1,59_ = 0.412, p = 0.523, η2 = 0.00683, 95% CI[-0.303, 0.156]; Interaction: F_1,59_ = 0.895, p = 0.348, η2 = 0.0148, 95% CI[-0.676, 0.242]). We also analyzed the mEPSC amplitude as a measure of postsynaptic AMPAR drive and found no significant genotype differences in either D1-SPNs or D2-SPNs (Figures 1E-F, Genotype: F_1,59_ = 1.35, p = 0.249, η2 = 0.0221, 95% CI[-1.97, 0.521]; SPN subtype: F_1,59_ = 0.431, p = 0.514, η2 = 0.00704, 95% CI[-0.837, 1.65]; Interaction: F_1,59_ = 0.504, p = 0.48, η2 = 0.00823, 95% CI[-3.37, 1.61]). These results indicate that *Fmr1* deletion does not impair the number of glutamatergic synapses or postsynaptic AMPAR function onto DMS D1- SPNs and D2-SPNs at P14-15, consistent with normal postnatal development of glutamatergic synaptic input.

**Figure 1.**
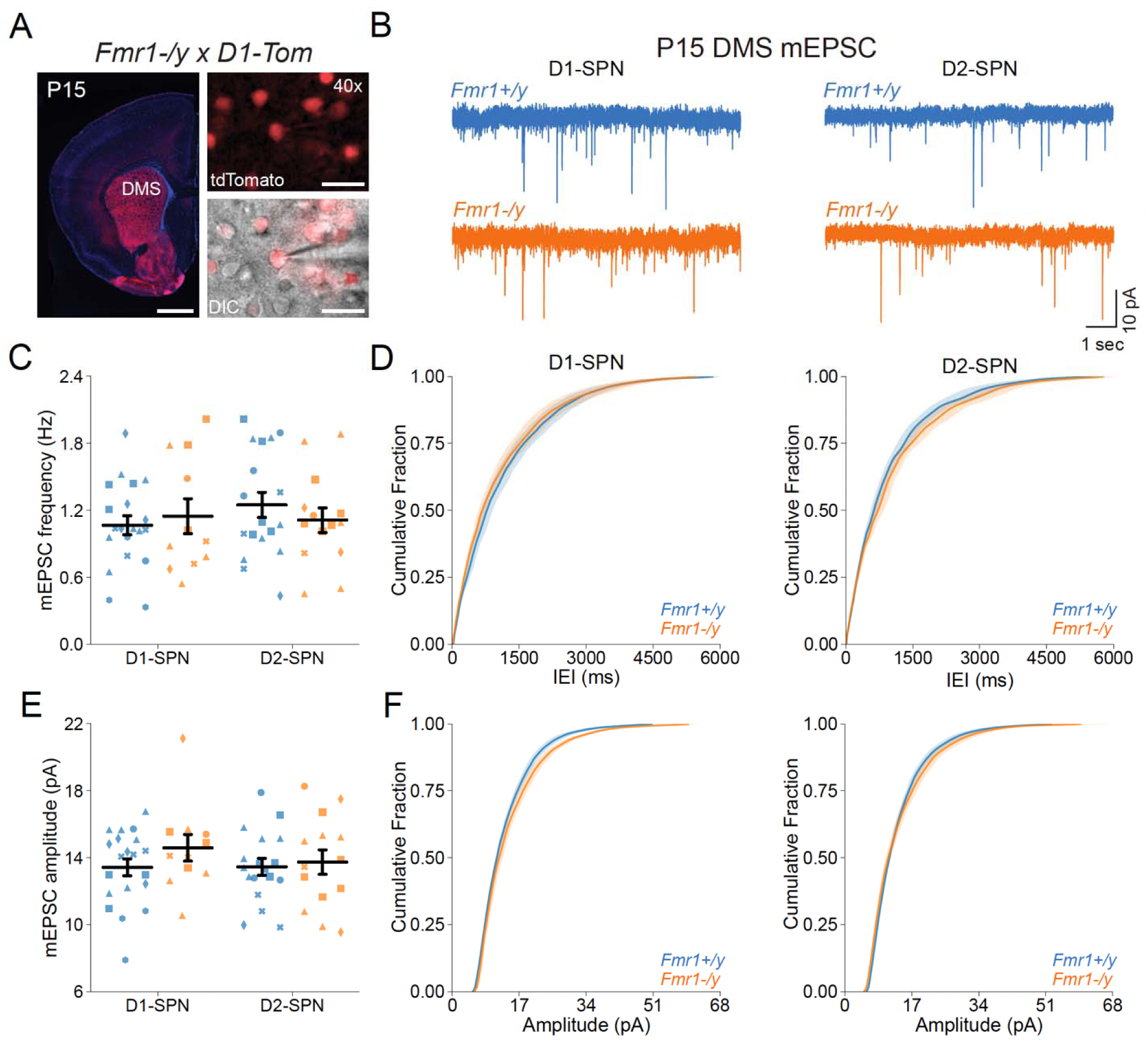
Normal glutamatergic synaptic transmission in D1- and D2-SPNs of the DMS in P14-15 *Fmr1*-/y mice. **(A)** Schematic representing a coronal brain section and whole-cell recordings of D1-SPNs (labeled with D1-Tomato) and D2-SPNs (td-Tomato negative) in the DMS of P14-15 *Fmr1*-/y and *Fmr1+/y* mice. **(B)** Representative AMPAR-mediated mEPSCs in D1-SPNs (left) and D2-SPNs (right) of *Fmr1+/y* (blue) and *Fmr1*-/y (orange) mice. **(C)** Average mEPSC frequency. **(D)** Cumulative distribution of inter-event intervals (IEIs) of mEPSCs in D1- and D2-SPNs (left and right, respectively). **(E)** Average mEPSC amplitude. **(F)** Cumulative distribution of mEPSC amplitudes in D1- and D2-SPNs (left and right respectively). Plots show individual data points, with shape representing a mouse within the group (i.e. Fmr1-/y x D1- SPN) and the summary with mean ± SEM. *P<0.05, **P<0.01, ***P<0.005, ****P<0.001.

### Normal Intrinsic excitability and passive membrane properties of DMS D1- and D2-SPNs of *Fmr1*-/y mice at P15

DMS SPNs undergo extensive maturation of their intrinsic properties during postnatal development ^39,47^. To determine whether *Fmr1* deletion impacts the excitability and passive membrane properties of D1- and D2-SPNs at P15, we performed current-clamp whole-cell recordings in acute brain slices from postnatal *Fmr1*-/y and *Fmr1+/y* mice. Recordings were performed at near-physiological temperature using a potassium-based internal solution to measure membrane voltage changes in response to stepped current injections (Figure 2A). We observed no genotype differences in the relationship between injected current amplitude and action potential (AP) firing (I-F) in both D1- and D2-SPNs (Figures 2B; Genotype: F_1,41_ = 1.47, p = 0.233; SPN subtype: F_1,41_ = 24.84, p<0.001; Genotype*SPN Subtype: F_1,41_ = 0.04, p=0.848; Pulse Amplitude: F_11,451_ = 362.53, p <0.001; Genotype*Pulse Amplitude: F_11,451_ = 1.35, p=0.192; SPN subtype*Pulse Amplitude: F_11,451_ = 18.66, p<0.001; Genotype*SPN subtype*Pulse amplitude: F_11,451_ = 0.09, p = 1.00). Quantification of membrane resistance also revealed no significant differences between genotypes for either D1- or D2-SPNs (Figure 2C, Genotype: F_1,41_ = 2.62, p = 0.113, η2 = 0.0467, 95% CI[-49.1, 5.4]; SPN subtype: F_1,41_ = 12.5, p = 0.00103, η2 = 0.223, 95% CI[-75.0, -20.5]; Interaction: F_1,41_ = 0.00805, p = 0.929, η2 = 0.000143, 95% CI[-52.1, 56.9]). Similarly, resting membrane potential (RMP) was unaffected by genotype (Figure 2D; Genotype: F_1,41_ = 0.257, p = 0.615, η2 = 0.0055, 95% CI[-3.9, 2.33]; SPN subtype: F_1,41_ = 5.49, p = 0.0241, η2 = 0.117, 95% CI[-6.73, -0.498]; Interaction: F_1,41_ = 0.0649, p = 0.8, η2 = 0.00139, 95% CI[-5.45, 7.02]). We next assessed rheobase current, an indicator of neuronal excitability, and found no significant differences between genotypes (Figure 2E; Genotype: F_1,41_ = 0.501, p = 0.483, η2 = 0.0086, 95% CI[-10.1, 20.9]; SPN subtype: F_1,41_ = 16.6,p = 0.000203, η2 = 0.286, 95% CI[15.8, 46.8]; Interaction: F_1,41_ = 0.0651, p = 0.8, η2 = 0.00112, 95% CI[-27.1, 34.9]). These findings indicate that *Fmr1* deletion does not significantly alter the intrinsic properties or passive membrane characteristics of D1- or D2-SPNs in the DMS at P14- 15.

**Figure 2.**
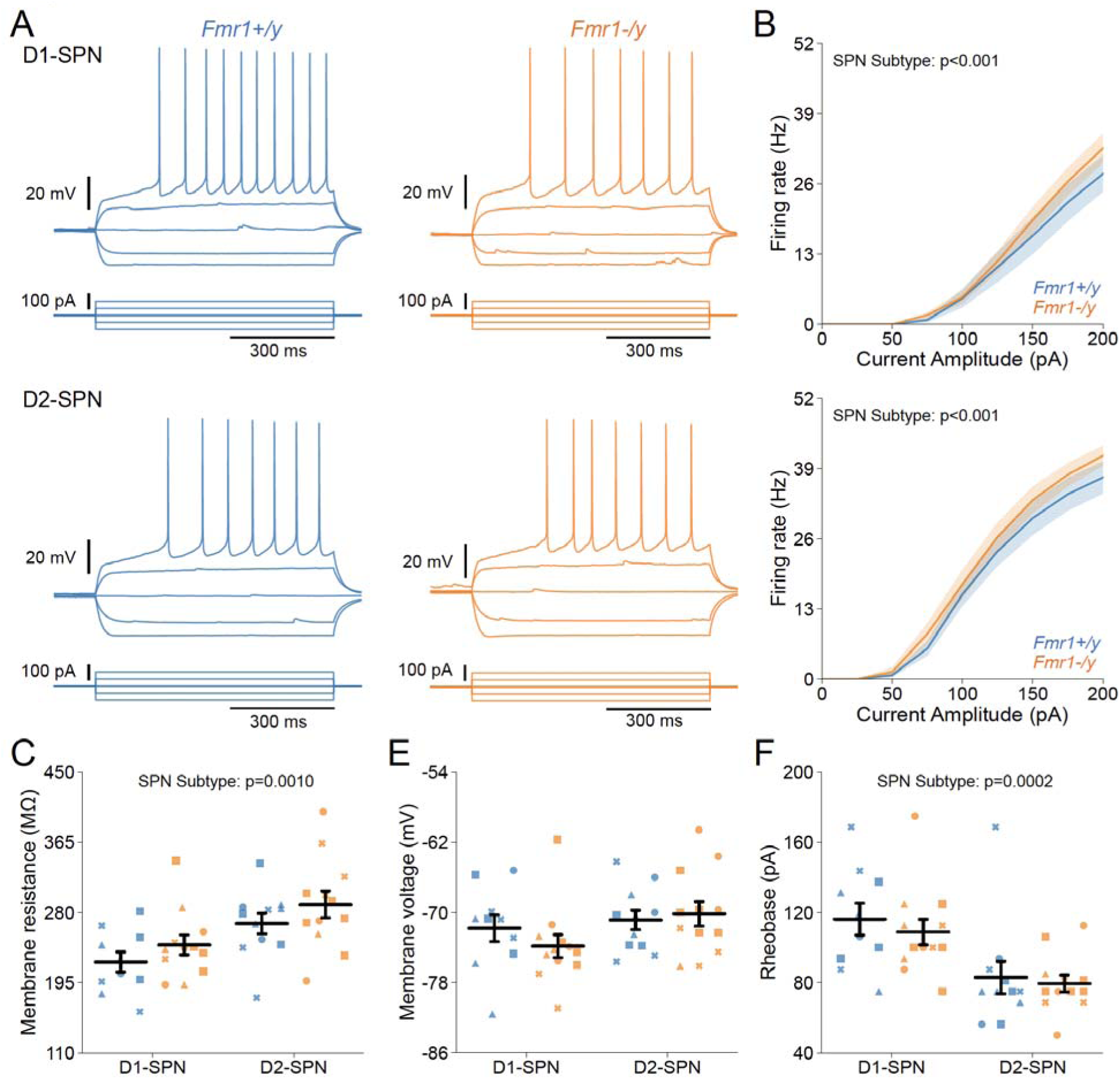
Normal intrinsic excitability and passive membrane properties of D1- and D2- SPNs in P14-15 *Fmr1*-/y mice. (A) Representative current-clamp recordings showing action potential firing in response to current injections in D1-SPNs (top) and D2-SPNs (bottom) of *Fmr1-/y* (orange) and *Fmr1+/y* (blue) P-15 mice. **(B)** Input-output relationship of firing frequency versus current amplitude (I-F curves) in D1- and D2-SPNs (top and bottom respectively). **(C)** Average membrane resistance in D1- and D2-SPNs. **(D)** Average resting membrane potential in D1- and D2-SPNs. **(E)** Average rheobase current in D1- and D2-SPNs. Plots show individual data points, with shape representing a mouse within the group (i.e. Fmr1-/y x D1-SPN) and the summary with mean ± SEM. *P<0.05, **P<0.01, ***P<0.005, ****P<0.001.

### Normal action potential properties of DMS D1- and D2-SPNs of *Fmr*1-/y mice at P15

Deletion of *Fmr1* alters action potential (AP) kinetics of adult NAc SPNs ^36^. To investigate whether Fmr1 deletion similarly affects APs of DMS D1- and D2-SPNs during postnatal development, we analyzed AP waveforms from rheobase traces obtained during our current-clamp recordings (Figures 3A and D). AP threshold was unaffected by genotype in both D1- SPNs and D2-SPNs (Figure 3B, Genotype: F_1,41_ = 1.27, p = 0.266, η2 = 0.0284, 95% CI[-1.77, 0.5]; SPN subtype: F_1,41_ = 2.54, p = 0.119, η2 = 0.0565, 95% CI[-0.24, 2.03]; Interaction: F_1,41_ = 0.066, p = 0.799, η2 = 0.00147, 95% CI[-2.55, 1.98]). Quantification of AP half-width revealed no significant differences between genotypes for either D1- or D2-SPNs (Figure 3C; Genotype: F_1,41_ = 0.543, p = 0.465, η2 = 0.011, 95% CI[-0.0357, 0.0767]; SPN subtype: F_1,41_ = 7.52, p = 0.009, η2 = 0.153, 95% CI[-0.132, -0.0201]; Interaction: F_1,41_ = 0.216, p = 0.644, η2 = 0.00439, 95% CI[-0.0865, 0.138]). Additionally, we found no differences in the peak AP velocity (Figure 3E; Genotype: F_1,41_ = 0.0022, p = 0.963, η2 = 5.06e-05, 95% CI[-1.51, 1.59]; SPN subtype: F_1,41_ = 2.19, p = 0.146, η2 = 0.0503, 95% CI[-0.413, 2.69]; Interaction: F_1,41_ = 0.386, p = 0.538, η2 = 0.00885, 95% CI[-4.05, 2.15]) or the time at which peak AP velocity occurred (Figure S1A; Genotype: F_1,41_ = 1.67, p = 0.203, η2 = 0.0372, 95% CI[-0.00812, 0.0371]; SPN subtype: F_1,41_ = 0.321, p = 0.574, η2 = 0.00713, 95% CI[-0.0163, 0.029]; Interaction: F_1,41_ = 2.02, p = 0.162, η2 = 0.045, 95% CI[-0.0134, 0.0771]). We also found no differences in the minimum AP velocity (Figure 3F; Genotype: F_1,41_ = 0.0531, p = 0.819, η2 = 0.000983, 95% CI[-0.621, 0.494]; SPN subtype: F_1,41_ = 12.7, p = 0.000957, η2 = 0.235, 95% CI[-1.54, -0.425]; Interaction: F_1,41_ = 0.266, p = 0.609, η2 = 0.00493, 95% CI[-0.83, 1.4]) or time of minimum AP velocity (Figure S1B; Genotype: F_1,41_ = 0.695, p = 0.409, η2 = 0.0158, 95% CI[-0.0332, 0.0798]; SPN subtype: F_1,41_ = 1.21, p = 0.277, η2 = 0.0276, 95% CI[-0.0873, 0.0257]; Interaction: F_1,41_ = 1.15, p = 0.29, η2 = 0.0261, 95% CI[-0.053, 0.173]). Interestingly, peak action potential voltage was significantly lower in D1-SPNs of Fmr1-/y mice, the only significantly different SPN physiological property we detected at this age (Figure S1C; Genotype: F_1,41_ = 6.54, p = 0.0144, η2 = 0.121, 95% CI[0.531, 4.53]; SPN subtype: F_1,41_ = 5.47, p = 0.0243, η2 = 0.101, 95% CI[0.316, 4.31]; Interaction: F_1,41_ = 1.18, p = 0.284, η2 = 0.0218, 95% CI[-1.85, 6.14]). Finally, we evaluated the peak afterhyperpolarization (AHP) amplitude and found no significant differences between genotypes (Figure S1D; Genotype: F_1,41_ = 0.594, p = 0.445, η2 = 0.0141, 95% CI[-2.0, 0.894]; SPN subtype: F_1,41_ = 0.382, p = 0.54, η2 = 0.00906, 95% CI[-1.89, 1.0]; Interaction: F_1,41_ = 0.195, p = 0.661, η2 = 0.00461, 95% CI[-3.52, 2.26]) or the time of peak AHP (Fig S1E; Genotype: F_1,41_ = 0.981, p = 0.328, η2 = 0.0187, 95% CI[-0.0496, 0.145]; SPN subtype: F_1,41_ = 9.44, p = 0.00377, η2 = 0.18, 95% CI[-0.246, -0.0508]; Interaction: F_1,41_ = 0.94, p = 0.338, η2 = 0.018, 95% CI[-0.101, 0.288]). These results indicate that *Fmr1* deletion does not significantly affect the AP waveform of DMS SPNs at P14-15, with the exception of a modest decrease in AP peak voltage in D1-SPNs.

**Figure 3.**
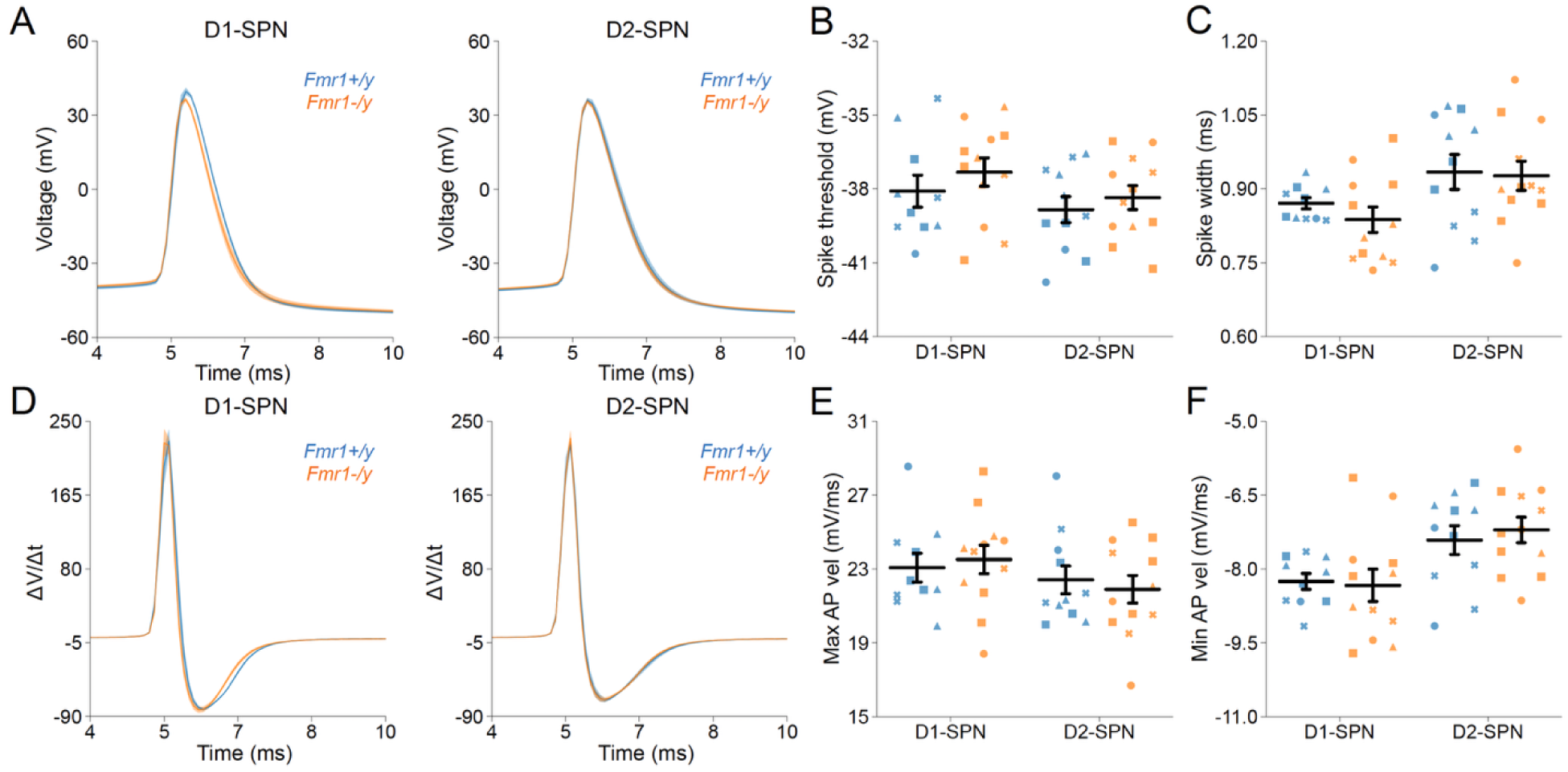
Action potential properties in D1- and D2-SPNs of P14-15 *Fmr1*-/y mice. **(A)** Representative single action potential traces recorded from D1-SPNs (left) and D2-SPNs (right) of *Fmr1-/y* (orange) and *Fmr1+/y* (blue) P-15 mice. **(B)** Average AP threshold in D1- and D2- SPNs. **(C)** Average AP half-width in D1- and D2-SPNs. **(D)** AP velocity (ΔV/Δt) in D1- and D2- SPNs (left and right respectively). **(E)** Maximum AP velocity. **(F)** Minimum AP velocity. Plots show individual data points, with shape representing a mouse within the group (i.e. Fmr1-/y x D1-SPN) and the summary with mean ± SEM. *P<0.05, **P<0.01, ***P<0.005, ****P<0.001.

### Normal glutamatergic synaptic input in DMS D1- and D2-SPNs of adult *Fmr1*-/y mice

Previous studies have shown region-specific changes in dendritic spine density and morphology in SPNs of the NAc and DLS in adult *Fmr1*-/y mice ^13,15^. Given our findings during postnatal development, where no synaptic deficits were detected (Figure 1), we sought to determine whether *Fmr1* loss leads to disruptions in glutamatergic inputs to DMS D1- and D2-SPNs later in development. To address this, we recorded AMPAR-mediated mEPSCs in acute brain slices of adult P60 *Fmr1*-/y and Fmr1+/y mice (Figure 4A). Quantification of mEPSC frequency revealed no significant difference between genotypes in either D1-SPNs or D2-SPNs (Figure 4C-D; Genotype: F_1,101_ = 0.749, p = 0.389, η2 = 0.00598, 95% CI[-0.591, 0.232]; SPN subtype:

**Figure 4.**
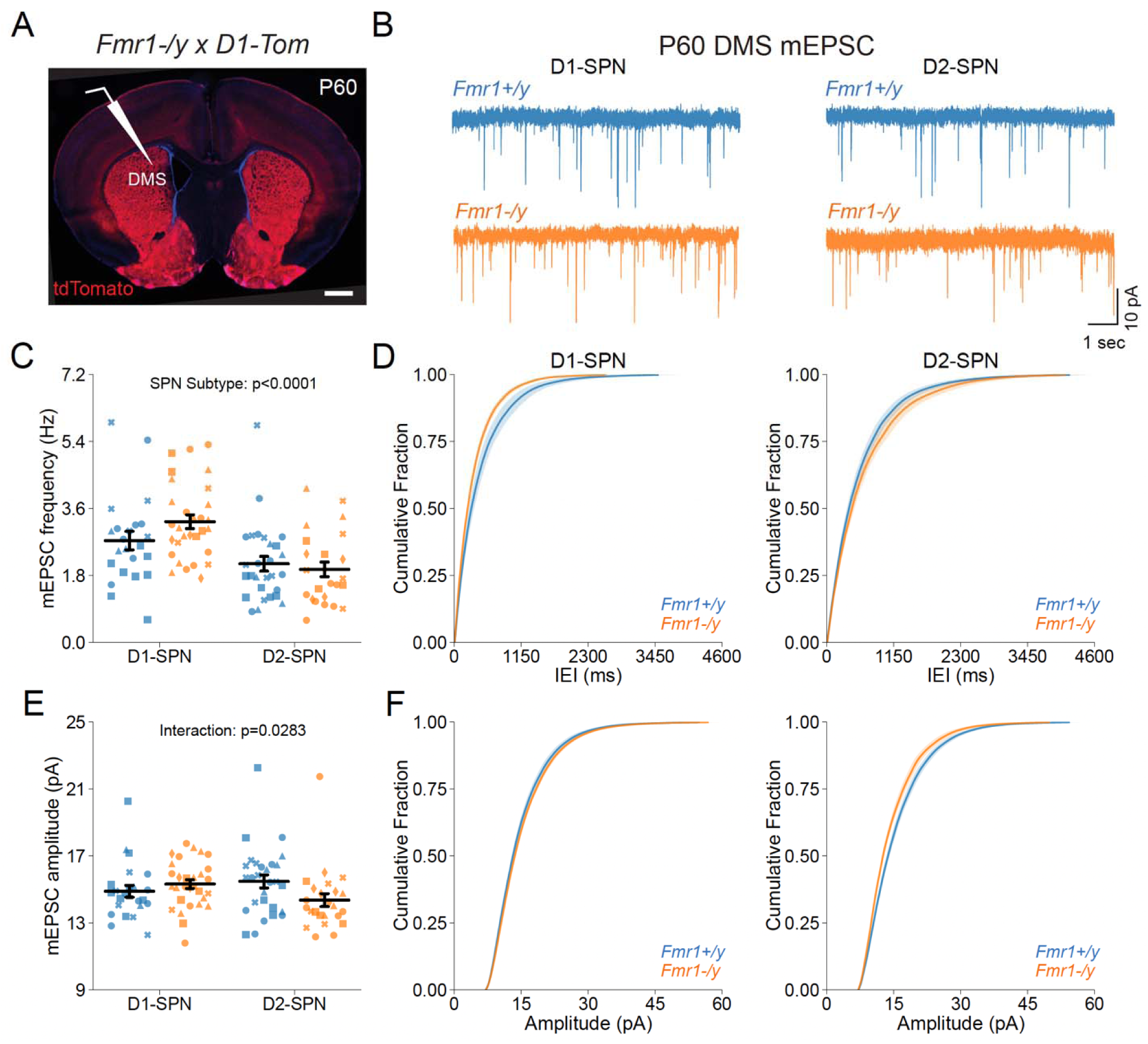
Normal glutamatergic synaptic transmission in D1- and D2-SPNs of the DMS in adult (P60) *Fmr1*-/y mice. **(A)** Schematic representing a coronal brain section and whole-cell recordings of D1-SPNs (labeled with D1-Tomato) and D2-SPNs in the DMS of P60 *Fmr1*-/y and *Fmr1+/y* mice. **(B)** Representative AMPAR-mediated mEPSCs in D1-SPNs (left) and D2-SPNs (right) of *Fmr1-/y* (orange) and *Fmr1+/y* (blue) mice. **(C)** Average mEPSC frequency . **(D)** Cumulative distribution of inter-event intervals (IEIs) of mEPSCs in D1- and D2-SPNs (left and right respectively). **(E)** Average mEPSC amplitude. **(F)** Cumulative distribution of mEPSC amplitudes in D1- and D2-SPNs (left and right, respectively). Plots show individual data points with shape representing a mouse within the group (i.e. Fmr1-/y x D1-SPN) and the summary with mean ± SEM. *P<0.05, **P<0.01, ***P<0.005, ****P<0.001.

F_1,101_ = 21.1, p = 1.26e-05, η2 = 0.168, 95% CI[0.541, 1.36]; Interaction: F_1,101_ = 2.51, p = 0.116, η2 = 0.0201, 95% CI[-1.48, 0.165]). We also analyzed mEPSC amplitude and found no significant interaction between genotype and SPN subtype (Figure 4E-F; Genotype: F_1,101_ = 0.985, p = 0.323, η2 = 0.00918, 95% CI[-0.345, 1.04]; SPN subtype: F_1,101_ = 0.275, p = 0.601, η2 = 0.00257, 95% CI[-0.508, 0.874]; Interaction: F_1,101_ = 4.95, p = 0.0283, η2 = 0.0462, 95% CI[-2.93, -0.168]). Taken together, these findings suggest that *Fmr1* deletion does not affect the number or strength of glutamatergic synapses onto D1-SPNs and D2-SPNs in the DMS during postnatal development or early adulthood.

### Hyperexcitability of DMS D1- and D2-SPNs in P60 *Fmr1*-/y mice

We further characterized the intrinsic properties of DMS SPNs in adult *Fmr1*-/y mice by performing whole-cell current-clamp recordings, as previously described (Figure 2). Notably, we detected a pronounced increase in the I-F relationship in both D1- and D2-SPNs of *Fmr1*-/y mice, with a more severe effect observed in D1-SPNs (Figures. 5B; Genotype: F_1,50_ = 6.55, p = 0.0135; SPN subtype: F_1,50_ = 22.98, p<0.001; Genotype*SPN Subtype: F_1,50_ = 1.40, p=0.0119; Pulse Amplitude: F_12,600_ = 548, p =0.863; Genotype*Pulse Amplitude: F_12,600_ = 5.16, p=0.0558; SPN subtype*Pulse Amplitude: F_12,600_ = 21.6, p=0.198; Genotype*SPN subtype*Pulse amplitude: F_12,600_ = 1.82, p = 0.0204). We also observed elevated membrane resistance in both *Fmr1*-/y D1- and D2-SPNs compared to Fmr1+/y controls (Figure 5C; Genotype: F_1,77_ = 5.0, p = 0.0283, η2 = 0.0544, 95% CI[-32.2, -1.86]; SPN subtype: F_1,77_ = 9.07, p = 0.00351, η2 = 0.0989, 95% CI[-38.2, -7.79]; Interaction: F_1,77_ = 0.698, p = 0.406, η2 = 0.0076, 95% CI[-43.1, 17.6]), but no genotype difference in resting membrane potential (Figure 5D; Genotype: F_1,76_ = 1.43, p = 0.235, η2 = 0.0176, 95% CI[-4.5, 1.12]; SPN subtype: F_1,76_ = 3.26, p = 0.0751, η2 = 0.0401, 95% CI[-5.35, 0.264]; Interaction: F_1,76_ = 0.581, p = 0.448, η2 = 0.00715, 95% CI[-7.77, 3.47]).

**Figure 5.**
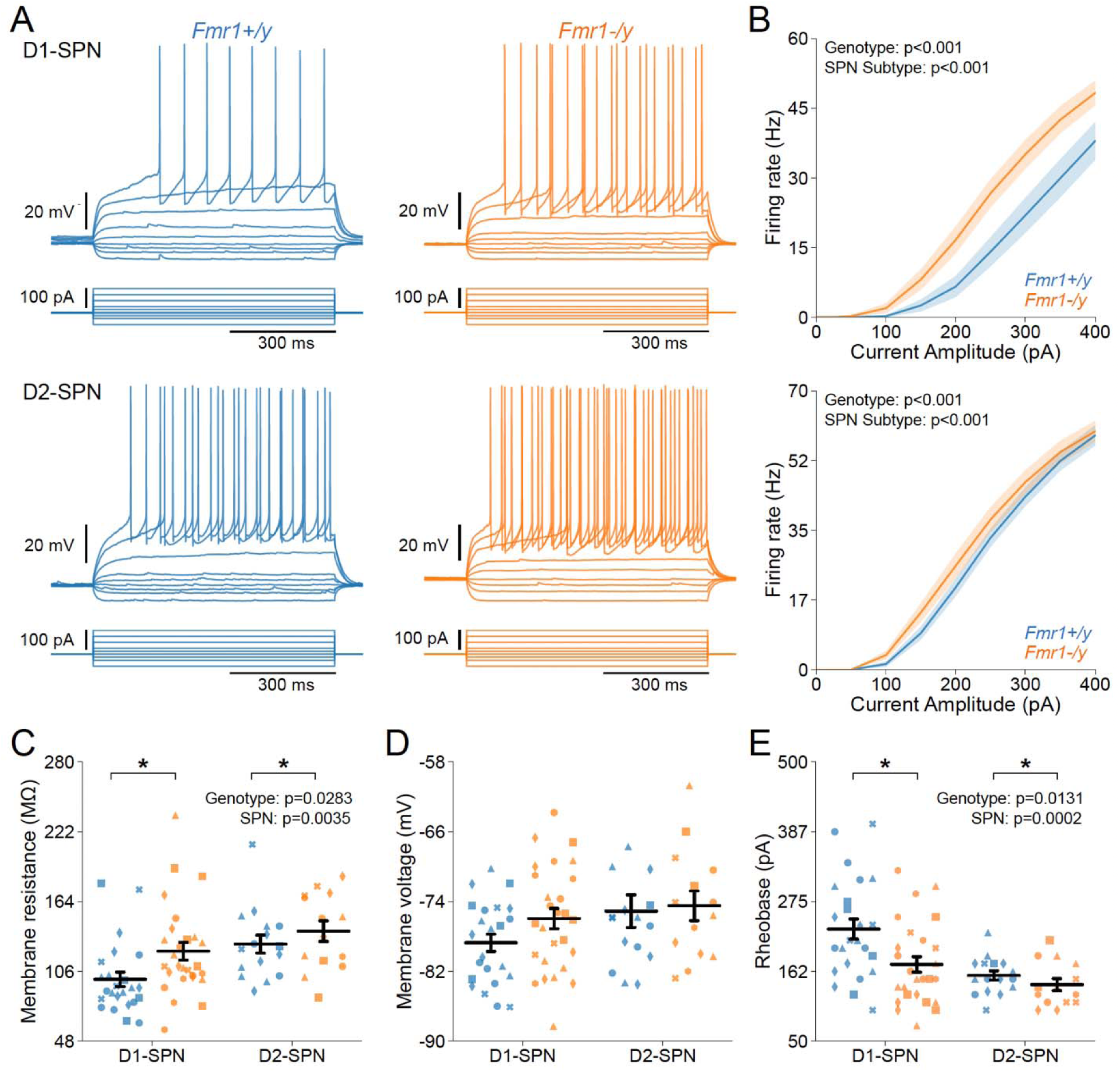
increased intrinsic excitability of D1- and D2-SPNs in adult P60 *Fmr1*-/y mice. **(A)** Representative current-clamp recordings showing action potential firing in response to current injections in D1-SPNs (top) and D2-SPNs (bottom) of *Fmr1+/y* (blue) and *Fmr1*-/y (orange) P60 mice. **(B)** Input-output relationship of firing frequency versus current amplitude (I-F curves) in D1- and D2-SPNs (top and bottom respectively). **(C)** Average membrane resistance. **(D)** Average resting membrane potential. **(E)** Average rheobase current. Plots show individual data points, with shape representing a mouse within the group (i.e. Fmr1-/y x D1-SPN) and the summary with mean ± SEM. *P<0.05, **P<0.01, ***P<0.005, ****P<0.001.

Consistent with neural hyperexcitability and I-F relationship, *Fmr1*-/y D1- and D2-SPNs exhibited lower rheobase currents compared to Fmr1+/y (Figure 5E; Genotype: F_1,77_ = 6.45, p = 0.0131, η2 = 0.0644, 95% CI[7.73, 63.9]; SPN subtype: F_1,77_ = 14.5, p = 0.000284, η2 = 0.144, 95% CI[25.6, 81.7]; Interaction: F_1,77_ = 2.24, p = 0.139, η2 = 0.0224, 95% CI[-14.0, 98.4]). These results indicate that *Fmr1* deletion causes a substantial increase in D1-SPN and D2-SPN excitability in the DMS, that emerges later in development.

### Altered action potential properties of DMS D1- and D2-SPNs in P60 *Fmr1*-/y mice

To determine whether changes in membrane resistance and rheobase observed in P60 *Fmr1*-/y SPNs are associated with abnormal AP properties, we analyzed AP parameters from whole-cell current-clamp recordings (Figures 6A and D). Analysis of AP onset revealed lower AP threshold in *Fmr1*-/y D1-SPNs compared to Fmr1+/y, while D2-SPNs showed a small increase (Figure 6B; Genotype: F_1,77_ = 2.03, p = 0.159, η2 = 0.0244, 95% CI[-0.732, 4.4]; SPN subtype: F_1,77_ = 0.366, p = 0.547, η2 = 0.00441, 95% CI[-1.79, 3.35]; Interaction: F_1,77_ = 3.65, p = 0.0599, η2 = 0.0439, 95% CI[-0.211, 10.1]). Quantification of AP half-width revealed broader APs in *Fmr1*-/y D1-SPNs compared to Fmr1+/y, with a more modest increase observed in D2-SPNs (Figure 6C; Genotype: F_1,77_ = 11.8, p = 0.000936, η2 = 0.13, 95% CI[-0.114, -0.0304]; SPN subtype: F_1,77_ = 0.967, p = 0.329, η2 = 0.0106, 95% CI[-0.0212, 0.0624]; Interaction: F_1,77_ = 1.14, p = 0.29, η2 = 0.0125, 95% CI[-0.128, 0.0388]). We next analyzed peak AP velocity and found no significant differences between genotypes (Figure S2A; Genotype: F_1,77_ = 0.0961, p = 0.757, η2 = 0.00122, 95% CI[-1.31, 1.8]; SPN subtype: F_1,77_ = 1.67, p = 0.2, η2 = 0.0212, 95% CI[-2.56, 0.545]; Interaction: F_1,77_ = 0.19, p = 0.664, η2 = 0.0024, 95% CI[-3.79, 2.43]) but observed delayed time of peak AP velocity (Figure S2B; Genotype: F_1,77_ = 4.39, p = 0.0395, η2 = 0.0513, 95% CI[- 0.0318, -0.0008]; SPN subtype: F_1,77_ = 2.05, p = 0.156, η2 = 0.024, 95% CI[-0.00435, 0.0266]; Interaction: F_1,77_ = 2.12, p = 0.149, η2 = 0.0248, 95% CI[-0.0537, 0.00831]). Likewise, the minimum AP velocity was decreased (Figure 6E; Genotype: F_1,77_ = 3.81, p = 0.0545, η2 = 0.047, 95% CI[-1.84, 0.0181]; SPN subtype: F_1,77_ = 0.00762, p = 0.931, η2 = 9.4e-05, 95% CI[-0.889, 0.971]; Interaction: F_1,77_ = 0.287, p = 0.594, η2 = 0.00353, 95% CI[-2.36, 1.36]) and the time of minimum AP velocity was also delayed (Fig 6F; Genotype: F_1,77_ = 13.0, p = 0.00055, η2 = 0.139, 95% CI[-0.126, -0.0364]; SPN subtype: F_1,77_ = 1.73, p = 0.192, η2 = 0.0186, 95% CI[-0.0152, 0.0746]; Interaction: F_1,77_ = 1.74, p = 0.191, η2 = 0.0186, 95% CI[-0.149, 0.0304]). AP peak voltage was not changed (Figure S2C; Genotype: F_1,77_ = 2.63, p = 0.109, η2 = 0.0319, 95% CI[- 4.01, 0.41]; SPN subtype: F_1,77_ = 2.68, p = 0.106, η2 = 0.0325, 95% CI[-0.394, 4.03]; Interaction:

**Figure 6.**
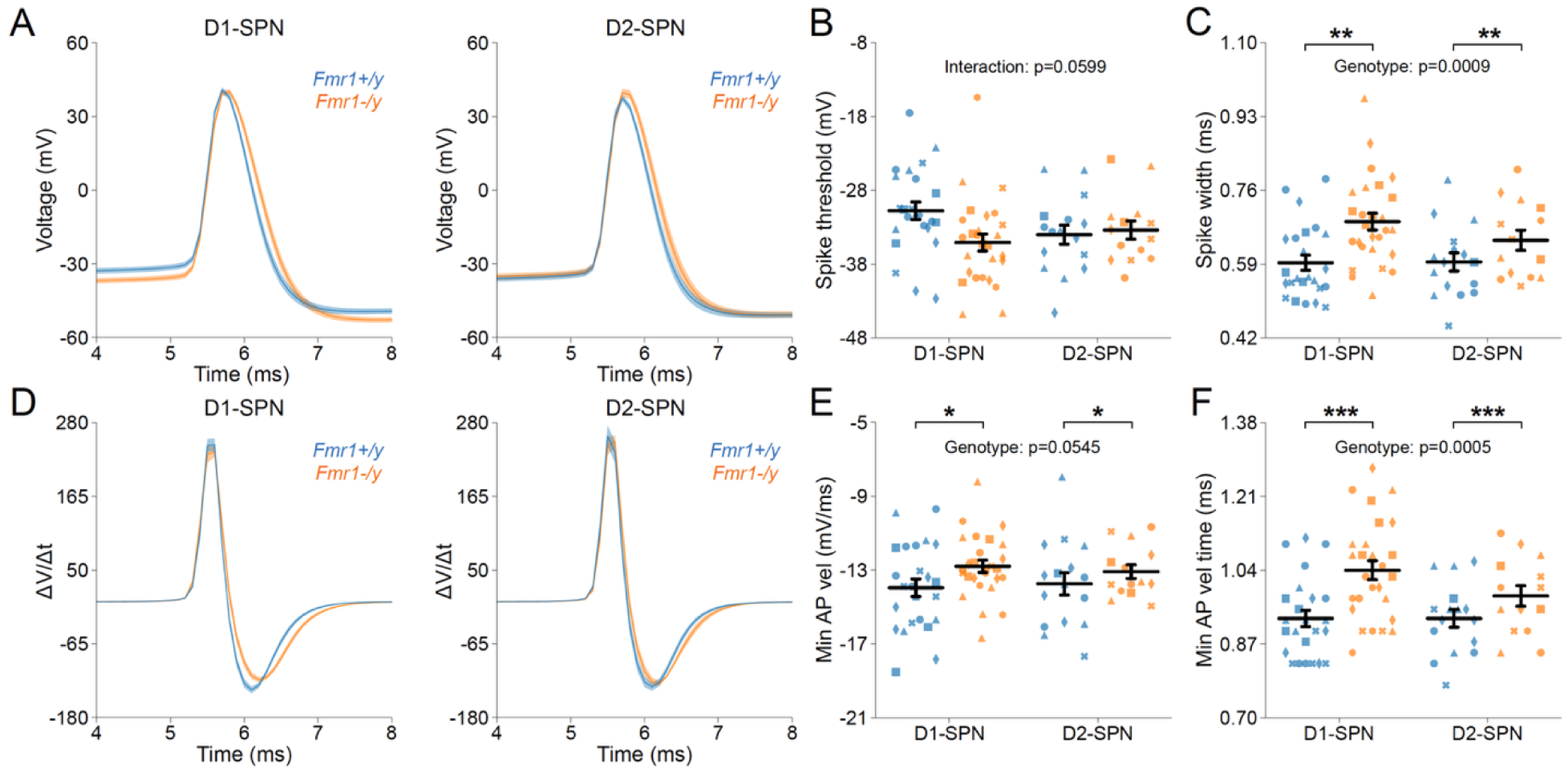
Action potential properties in D1- and D2-SPNs of adult P60 *Fmr1*-/y mice. **(A)** Representative single action potential traces recorded from D1-SPNs (left) and D2-SPNs (right) of *Fmr1-/y* (orange) and *Fmr1+/y* (blue) P60 mice. **(B)** Average AP half-width in D1- and D2- SPNs **(C)** AP threshold potential **(D)** AP velocity (ΔV/Δt) (left and right respectively). **(E)** Minimum AP velocity. **(F)** Time of the minimum AP velocity. Plots show individual data points, with shape representing a mouse within the group (i.e. Fmr1-/y x D1-SPN) and the summary with mean ± SEM. *P<0.05, **P<0.01, ***P<0.005, ****P<0.001.

F_1,77_ = 0.188, p = 0.666, η2 = 0.00228, 95% CI[-3.46, 5.38]) and peak AHP amplitude was also not altered (Figure S2D; Genotype: F_1,77_ = 1.39, p = 0.241, η2 = 0.0173, 95% CI[-1.05, 4.09]; SPN subtype: F_1,77_ = 0.107, p = 0.744, η2 = 0.00133, 95% CI[-2.99, 2.15]; Interaction: F_1,77_ = 1.97, p = 0.165, η2 = 0.0245, 95% CI[-1.52, 8.76]). The time of peak AHP was slightly delayed but did not reach statistical significance (Figure S2E; Genotype: F_1,77_ = 3.36, p = 0.0706, η2 = 0.0414, 95% CI[-2.95, 71.6]; SPN subtype: F_1,77_ = 0.69, p = 0.409, η2 = 0.00849, 95% CI[-21.7, 52.8]; Interaction: F_1,77_ = 0.162, p = 0.689, η2 = 0.00199, 95% CI[-89.6, 59.5]). These findings indicate that *Fmr1* deletion alters specific AP properties in both D1-SPN and D2-SPN, including AP half-width, threshold, and peak AHP time.

### Chronic aripiprazole treatment does not normalize DMS SPN hyperexcitability in *Fmr1*-/y mice

To determine whether chronic aripiprazole treatment rescues the hyperexcitability phenotype observed in DMS SPNs in *Fmr1*-/y mice, we performed whole-cell current-clamp recordings in acute brain slices of adult *Fmr1*-/y and Fmr1+/y mice treated daily with either aripiprazole (3 mg/kg) or saline for 14 days (Figure 7A). As previously observed, SPNs of *Fmr1*-/y are hyperexcitable when compared to WT SPNs (Figure 7B). Aripiprazole treatment had no effect in this genotype difference and the I-F relationship was unaltered and remained elevated in *Fmr1*-/y SPNs compared to *Fmr1+/y*. (Figure 7C; Genotype: F_1,99_ = 12.69, p < 0.001; Treatment: F_1,99_ = 1.2, p=0.276; Genotype*Treatment: F_1,99_ = 0.6, p=0.441; Pulse Amplitude: F_12,1188_ = 920, p < 0.001; Genotype*Pulse Amplitude: F_12,1188_ = 10.06, p < 0.001; Treatment*Pulse amplitude: F_12,1188_ = 1.07, p=0.382; Genotype*Treatment*Pulse amplitude: F_12,1188_ = 0.49, p = 0.921). Quantification of membrane resistance also confirmed our results described in Figure 5 and revealed a significant increase in membrane resistance in *Fmr1*-/y SPNs compared to Fmr1+/y, but no significant effect of aripiprazole treatment was observed in either group (Figure 7D, Genotype: F_1,101_ = 6.58, p = 0.0118, η2 = 0.0596, 95% CI[-28.8, -3.69]; Treatment: F_1,101_ = 1.93, p = 0.168, η2 = 0.0175, 95% CI[-3.77, 21.4]; Interaction: F_1,101_ = 0.889, p = 0.348, η2 = 0.00805, 95% CI[-37.1, 13.2]). Likewise, AP threshold was hyperpolarized in *Fmr1*-/y mice, but unaffected by treatment (Figure 7E, Genotype: F_1,101_ = 8.46, p = 0.00446, η2 = 0.0754, 95% CI[0.729, 3.86]; Treatment: F_1,101_ = 1.69, p = 0.197, η2 = 0.015, 95% CI[-2.59, 0.54]; Interaction:

**Figure 7.**
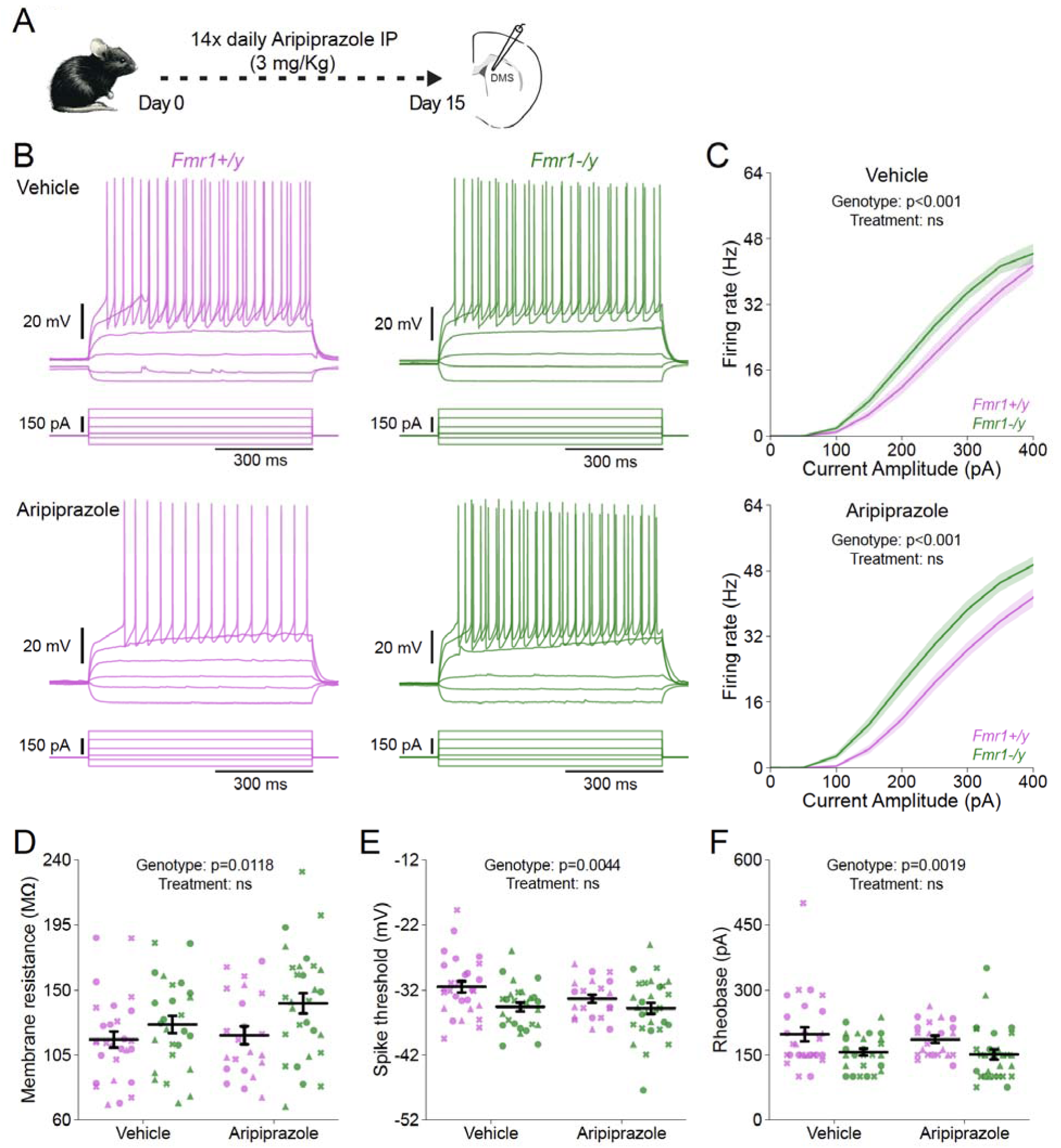
Chronic aripiprazole treatment does not ameliorate hyperexcitability in SPNs of *Fmr1*-/y mice. **(A)** Schematic showing daily intraperitoneal (IP) injections of aripiprazole (3 mg/kg) in adult mice for 14 days, followed by whole-cell recordings of SPNs in the DMS. (B) Representative current-clamp recordings showing action potential firing in response to current injections in D1-SPNs (top) and D2-SPNs (bottom) of Fmr1+/y (purple) and *Fmr1*-/y (green) P60 mice. **(C)** Input-output relationship of firing frequency versus current amplitude (I-F curves) in SPNs from *Fmr1*-/y and Fmr1+/y mice treated with vehicle (top) or aripiprazole (bottom). **(D)** Average membrane resistance. **(E)** Average AP threshold **(F)** Average rheobase current. Plots show individual data points, with shape representing a mouse within the group (i.e. Fmr1-/y x D1-SPN) and the summary with mean ± SEM. *P<0.05, **P<0.01, ***P<0.005, ****P<0.001.

F_1,101_ = 1.07, p = 0.303, η2 = 0.00957, 95% CI[-4.76, 1.49]), and the reduced rheobase in *Fmr1*-/y SPNs was also unaffected by aripiprazole treatment (Fig 7F, Genotype: F_1,101_ = 10.1, p = 0.00199, η2 = 0.0903, 95% CI[14.2, 61.5]; Treatment: F_1,101_ = 0.505, p = 0.479, η2 = 0.00452, 95% CI[-32.1, 15.2]; Interaction: F_1,101_ = 0.0827, p = 0.774, η2 = 0.00074, 95% CI[-54.1, 40.4]).

Some AP waveform properties (Figure S3A) such as peak AHP voltage were decreased in SPNs of *Fmr1-/y* mice compared to WT, but were unaffected by treatment (Figure S3B, Genotype: F_1,101_ = 5.57, p = 0.0202, η2 = 0.0513, 95% CI[0.308, 3.56]; Treatment: F_1,101_ = 0.258, p = 0.613, η2 = 0.00237, 95% CI[-2.04, 1.21]; Interaction: F_1,101_ = 1.75, p = 0.189, η2 = 0.0161, 95% CI[-5.42, 1.08]). Other AP properties showed no difference between genotypes when analyzed in combined SPN populations such as AP half-width (Figure S3C, Genotype: F_1,101_ = 0.942, p = 0.334, η2 = 0.00896, 95% CI[-0.055, 0.0189]; Treatment: F_1,101_ = 1.37, p = 0.245, η2 = 0.013, 95% CI[-0.0151, 0.0587]; Interaction: F_1,101_ = 1.87, p = 0.175, η2 = 0.0178, 95% CI[- 0.023, 0.125]), maximum AP velocity (Figures S3D and S3E, Genotype: F_1,101_ = 0.6, p = 0.441, η2 = 0.0059, 95% CI[-0.769, 1.75]; Treatment: F_1,101_ = 0.0074, p = 0.932, η2 = 7.28e-05, 95% CI[-1.32, 1.21]; Interaction: F_1,101_ = 0.0805, p = 0.777, η2 = 0.000791, 95% CI[-2.88, 2.16]), time of maximum AP velocity (Figure S3F, Genotype: F_1,101_ = 3.09, p = 0.0817, η2 = 0.0278, 95% CI[-0.0316, 0.0019]; Treatment: F_1,101_ = 3.25, p = 0.0743, η2 = 0.0293, 95% CI[-0.00152, 0.032]; Interaction: F_1,101_ = 3.67, p = 0.0581, η2 = 0.0331, 95% CI[-0.00113, 0.0658]), minimum AP velocity (Figure S3G, Genotype: F_1,101_ = 0.292, p = 0.59, η2 = 0.00281, 95% CI[-0.51, 0.891]; Treatment: F_1,101_ = 1.87, p = 0.175, η2 = 0.0179, 95% CI[-0.218, 1.18]; Interaction: F_1,101_ = 0.954, p = 0.331, η2 = 0.00917, 95% CI[-0.711, 2.09]), time of minimum AP velocity (Figure S3H, Genotype: F_1,101_ = 0.6, p = 0.441, η2 = 0.0059, 95% CI[-0.769, 1.75]; Treatment: F_1,101_ = 0.0074, p = 0.932, η2 = 7.28e-05, 95% CI[-1.32, 1.21]; Interaction: F_1,101_ = 0.0805, p = 0.777, η2 = 0.000791, 95% CI[-2.88, 2.16]), peak AP voltage (Figure S3I, Genotype: F_1,101_ = 0.738, p = 0.392, η2 = 0.00712, 95% CI[-0.962, 2.43]; Treatment: F_1,101_ = 0.87, p = 0.353, η2 = 0.00839, 95% CI[-2.5, 0.899]; Interaction: F_1,101_ = 0.998, p = 0.32, η2 = 0.00964, 95% CI[-1.68, 5.1]), resting membrane potential (Figure S3J, Genotype: F_1,101_ = 0.901, p = 0.345, η2 = 0.00879, 95% CI[-2.27, 0.801]; Treatment: F_1,101_ = 0.0856, p = 0.771, η2 = 0.000835, 95% CI[-1.31, 1.76]; Interaction: F_1,101_ = 0.502, p = 0.48, η2 = 0.0049, 95% CI[-4.17, 1.97]), and the time at which the peak AHP occurred (Figure S3K, Genotype: F_1,101_ = 0.405, p = 0.526, η2 = 0.00388, 95% CI[- 0.1, 0.0515]; Treatment: F_1,101_ = 1.14, p = 0.288, η2 = 0.0109, 95% CI[-0.035, 0.117]; Interaction: F_1,101_ = 1.87, p = 0.175, η2 = 0.0179, 95% CI[-0.0472, 0.256]). Together, these findings indicate that chronic aripiprazole treatment does not significantly alter the intrinsic or AP properties of DMS SPNs in either *Fmr1+/y* or *Fmr1-/y* mice.

## Discussion

Here, we characterized the developmental trajectory of SPN dysfunction in the DMS of *Fmr1*-/y mice, a widely used model of FXS. Our findings reveal a striking hyperexcitability of both D1- and D2-SPNs in the DMS of adult *Fmr1*-/y male mice (Figures 5 and 6). However, intrinsic and synaptic properties of both SPN populations were normal at P15 (Figures 1-3), indicating a delayed onset of SPN deficits in FXS that only manifest in later developmental stages. We found no significant differences in mEPSC frequency or amplitude in both D1- and D2-SPNs at P14-15 (Figure 1) or P60 (Figure 4), suggesting normal glutamatergic synaptic inputs in the DMS of *Fmr1*-/y mice. This aligns with previous observations of unaltered dendritic spine density in DMS SPNs ^15^. By contrast, D1-SPNs in the DLS of *Fmr1*-/y mice show increased dendritic spine density and elevated mEPSC frequency ^13^, suggesting region-specific abnormalities of SPN synaptic properties across the striatum.

Despite normal mEPSC properties, we observed pronounced hyperexcitability of D1-SPNs and D2-SPNs in the DMS of adult Fmr1-/y mice, characterized by increased membrane resistance and reduced rheobase current (Figure 5). We also found substantial changes in AP waveform properties including wider AP width and slower repolarization kinetics. Abnormalities in membrane excitability and AP kinetics have been observed in other neuron types of adult *Fmr1*-/y mice, including pyramidal neurons of the mPFC and hippocampus. Interestingly, these changes are cell type-specific and caused by distinct ion channel alterations. In the hippocampus, CA3 pyramidal neurons exhibit wider APs due to reduced BK channel function ^48^, whereas CA1 neurons show normal AP waveform abnormalities but lower input resistance caused by elevated expression of dendritic HCN channels ^49^. In the cortex, L2/3 pyramidal neurons exhibit shorter APs with faster decay, driven by enhanced A-type (Kv4) potassium currents ^50^, whereas extratelencephalic, but not intratelencephalic L5 pyramidal neurons exhibit a hyperpolarized AP threshold, caused by reduced HCN and Kv1 currents ^51^. These findings highlight complex, cell type-specific patterns of K+ channel dysfunction in *Fmr1*-/y mice, leading to distinct alterations in excitability and AP generation and kinetics across different brain regions. In DMS SPNs, the broader AP waveforms observed in our study suggests slower repolarization. Since BK channels facilitate rapid AP repolarization and Kv4 channels regulate subthreshold excitability, their dysfunction in SPN could cause slow AP decay without affecting threshold properties. Future studies identifying the precise channel alterations in SPNs will be critical for understanding striatal circuit dysfunction in *Fmr1*-/y mice.

Perhaps the most critical finding of this study is the late onset of SPN hyperexcitability in *Fmr1*-/y mice. At P14-15, SPNs exhibit normal intrinsic properties, with hyperexcitability only observed in adult stages (Figure 5). This delayed onset suggests that the increased excitability is unlikely to result solely from the absence of FMRP, because in rodents FMRP expression peaks in the striatum during perinatal periods ^38^. Instead, SPN hyperexcitability may arise as a secondary adaptation to other circuit disruptions induced by deletion of *Fmr1*. Notably, *Fmr1*-/y mice exhibit abnormal patterns of cortical activity from very early postnatal stages ^10–12^, suggesting that cortical dysfunction precedes the onset of SPN physiological deficits. Given the tight functional coupling between cortical and striatal circuits during postnatal development ^21,31,39^, abnormal patterns of cortical activity might progressively drive secondary adaptations in downstream striatal circuits, ultimately resulting in SPN hyperexcitability. Alternatively, intrinsic changes in SPNs might reflect abnormal plasticity mechanisms induced by striatal network disruptions, such as altered local interneuron function or dopaminergic signaling, which are known to be affected in *Fmr1*-/y mice ^11,12^.

The observed hyperexcitability of DMS SPNs in adult *Fmr1*-/y mice has significant implications for understanding motor and cognitive symptoms in FXS. The DMS is a key region for goal-directed behavior and cognitive flexibility, and imbalances in SPN activity have been linked to various neurodevelopmental disorders, including ASD ^20,21^. Notably, several core symptoms of FXS such as irritability, aggression, and self-injurious behaviors become more prevalent and severe with age ^52,53^. The delayed hyperexcitability of SPNs may be associated with the developmental trajectory of these behavioral symptoms, and future studies should explore the relationship between striatal activity deficits and the deterioration of maladaptive behaviors throughout development. Additionally, Fragile X-associated tremor/ataxia syndrome (FXTAS), a late-onset neurodegenerative disorder in *FMR1* premutation carriers, leads to progressive motor and cognitive decline in adulthood ^54^. The mechanisms driving FXTAS symptoms remain unclear, but our findings suggest that SPN deficits resulting from *Fmr1* deletion are not solely a consequence of loss of FMRP during early development. It is thus possible that the loss of SPN function in adulthood contributes to the motor impairments observed in FXTAS, raising the need for further investigation into the role of striatal dysfunction in FXTAS pathophysiology. Moreover, given the widespread use of aripiprazole to manage maladaptive behaviors in FXS, our findings also have therapeutic implications. As a partial agonist of dopamine D2 receptors, aripiprazole is often prescribed to address irritability and aggression in FXS ^55^. However, our data indicates that chronic aripiprazole treatment in adult mice does not rescue or meliorate any excitability phenotypes of DMS SPNs in *Fmr1-/y* mice (Figure 7). This suggests that aripiprazole’s therapeutic effects may not directly target the core mechanisms underlying D1- or D2-SPN hyperexcitability, highlighting the need for alternative therapeutic strategies aimed at correcting core striatal deficits in FXS. For instance, a positive allosteric modulator (PAM) of the muscarinic acetylcholine M4 receptor has recently been shown to correct synaptic plasticity deficits in SPNs of *Fmr1-/y* mice ^13^. However, whether M4R PAMs can also normalize SPN excitability differences remains untested.

While our study provides valuable insights into the developmental trajectory of SPN dysfunction in FXS, it also raises important questions that warrant further investigation. First, the molecular mechanisms underlying SPN hyperexcitability remain unclear. Identifying these mechanisms could offer new therapeutic targets for mitigating striatal dysfunction in FXS. Additionally, our analysis of synaptic function focused exclusively on AMPAR-mediated mEPSCs, leaving open the possibility that other synaptic properties may be affected. Investigating these aspects could provide a more comprehensive understanding of striatal connectivity deficits in FXS. Finally, the broader functional consequences of SPN hyperexcitability for striatal circuit dynamics and behavior remain unclear. Given the early cortical disruptions reported in *Fmr1*-/y mice, future studies should aim to determine how these cortical abnormalities impact downstream striatal circuits in vivo and how such changes contribute to deficits in goal-directed behavior, motor control, and cognitive flexibility. Addressing these questions will be critical for advancing our understanding of striatal contributions to the complex behavioral phenotype of FXS.

## Acknowledgements

We thank Susana da Silva for helpful comments on the manuscript. We thank Lan Chen for helping with whole-cell recordings and Andrew D’Agostino, Tasha Merchant and Sidney Dawkins for assistance with mouse husbandry and genotyping.

## Data Availability

No large datasets were generated or analyzed in this study. Electrophysiology data are backed up on a server managed by the Department of Psychiatry at the University of Pittsburgh and will be made available upon request.

## Author Contributions

R.T.P. and L.N. conceived and designed the study. L.N., M.J, M.M. and Y-C.S. performed whole-cell recordings. L.N. developed the scripts for electrophysiological data analysis and analyzed the data. All authors participated in data interpretation and discussion. L.N. and R.T.P. wrote the manuscript with input from all the other authors.

## Ethics declarations

### Ethics approval

All procedures in mice were approved by the Institutional Animal Care and Use Committee (IACUC) at the University of Pittsburgh in compliance with the guidelines described in the US National Institutes of Health *Guide for the Care and Use of Laboratory Animals*.

### Consent to Participate

Not applicable as only mice were used in this study.

## Methods

### Animals

All experimental manipulations on mice were performed in accordance with protocols approved by the Institutional Animal Use and Care Committee at the University of Pittsburgh in compliance with the guidelines described in the US National Institutes of Health *Guide for the Care and Use of Laboratory Animals*. Mice were housed on a 12/12hr light/dark cycle with chow and water provided ad libitum. Mice were weaned at P21-23 and separated by sex in cages of 2-5 animals of mixed genotypes. Fmr1 mutant mice B6.129P2-*Fmr1^tm1Cgr^*/J were obtained from The Jackson Laboratory (#003025). Characterization of neural properties by electrophysiology was performed in male *Fmr1+/y* and *Fmr1-/y* age-matched mice.

### Aripiprazole preparation and administration

Aripiprazole (Millipore-Sigma: 1042634) was dissolved at 10mg/mL in 100% DMSO and stored in the dark at room temperature. The Aripiprazole/DMSO solution was dissolved in 1% Tween- 80 in 0.9% (0.375 mg/mL final concentration) saline each day prior to injections. Vehicle solution contained DMSO (0.0375 mL/mL) and 1% Tween-80 in 0.9% saline. Mice were injected with 3 mg/kg of Aripiprazole or vehicle intraperitoneally. Mice were weighed on day 1 before the first injection and the injection volume was adjusted for weight. Adult mice were administered Aripiprazole for 14 days every afternoon. The day after the final injection (∼18 hrs later), tissue was collected for acute slice electrophysiology.

### Brain slice preparation and whole-cell electrophysiology

Acute brain slices were prepared following anesthesia by isoflurane inhalation and transcardiac perfusion with ice-cold artificial cerebrospinal fluid (ACSF) containing (in mM): 125 NaCl, 2.5 KCl, 25 NaHCO_3_, 2 CaCl_2_, 1 MgCl_2_, 1.25 NaH_2_PO_4_ and 25 glucose (310 mOsm per kg). Cerebral hemispheres were removed and transferred into a slicing chamber containing ice-cold ACSF. Coronal slices including ACC (275 µm thick) were cut with a Leica VT1200s vibratome and transferred for 10 min to a holding chamber containing choline-based solution consisting of (in mM): 110 choline chloride, 25 NaHCO_3_, 2.5 KCl, 7 MgCl_2_, 0.5 CaCl_2_, 1.25 NaH_2_PO_4_, 25 glucose, 11.6 ascorbic acid, and 3.1 pyruvic acid at 33°C. Slices were subsequently transferred to a chamber with pre-warmed ACSF (33°C) and gradually cooled down to room temperature (20–22°C). All recordings were obtained within 4 hours of slicing. Both ACSF and choline solution were constantly bubbled with 95% O_2_ and 5% CO_2_. Individual slices were transferred to a recording chamber mounted on an upright microscope (Olympus BX51WI) and continuously perfused (1–2 ml per minute) with ACSF at room temperature. Cells were visualized using a 40× water-immersion objective with infrared illumination. Whole-cell voltage clamp recordings were made from SPNs in the dorsomedial striatum. Recording electrode pipettes (3-4 MΩ) pulled from borosilicate glass (BF150-86-7.5, Sutter Instruments). Voltage-clamp recordings were performed with a Cs^+^-based internal solution containing (in mM): 130 CsMeSO_4_, 10 HEPES, 1.8 MgCl_2_, 4 Na_2_ATP, 0.3 NaGTP, and 8 Na_2_-phosphocreatine,10 CsCl_2,_ 3.3 QX-314 (Cl^−^ salt), (pH 7.3 adjusted with CsOH; 300 mOsm per kg). Alexa Fluor 488 dye (0.07%) was added to the internal solution to fill the patched cell and confirm spiny morphology of SPNs. In voltage-clamp experiments, errors due to voltage drop across the series resistance (<20 MΩ) were left uncompensated. For mEPSC recordings, ACSF contained 1 μM TTX, 1 μM (RS)-CPP, and 1 μM Gabazine, and recordings were performed at room temperature (20-22°C) with V_m_ = -70 mV. After breaking in, cells were left to stabilize for 4 min and currents were then acquired continuously for 5 min. Membrane currents and potentials were amplified and low-pass filtered at 3 kHz using Multiclamp 700B amplifier (Molecular Devices), digitized at 10 kHz, and acquired using National Instruments acquisition boards and a custom version of ScanImage written in MATLAB (Mathworks). Calculation of input resistance and membrane capacitance in voltage clamp recordings was performed by fitting evoked currents in response to -5 mV voltage steps in the first seconds after cell break-in. For current-clamp recordings, potassium-based internal solution containing (in mM): 130 KMeSO_3_, 10 HEPES, 3 KCl, 1 EGTA, 4 Na_2_ATP, 0.3 NaGTP, and 8 Na_2_-phosphocreatine, (pH 7.3 adjusted with KOH; 300 mOsm per kg) was used. No drug was added to the bath, and recordings were performed at 31-33°C. Cell was broken in with V_m_ = -70 mV and allowed to stabilize for 4 min in voltage-clamp. After stabilization, resting membrane potential was measured at I = 0. Current-clamp recording was performed by adjusting holding current to maintain the V_m_ at –75 mV. For each experiment four cycles of 300ms baseline, 700ms current injection step and a 2000 ms baseline ending the acquisition. . For the adult timepoints the following current steps were used: -100, -50, -25, 0, 25, 50, 100, 150, 200, 250, 300, 350, 400. For the P14 timepoint the following current steps were used: -100, -75, -50, -25, 0, 25, 50, 75, 100, 125, 150, 175, 200. For the aripiprazole experiment the following current steps were used: -100, -75, -50, -25, 0, 25, 50, 75, 100, 125, 150, 175, 200. Recording data was saved as Matlab files for subsequent off-line analysis.

### Electrophysiology data analysis

Adult D1/D2 mEPSCs were analyzed using a custom program in Igor Pro. For all other experiments mEPSCs and current-clamp traces were analyzed using a custom python-based program ClampSuite available at https://github.com/LarsHenrikNelson/ClampSuite. For mEPSC analysis, acquisition offset was removed by subtracting the mean. Recordings were filtered using a zero-phase Remez filter with a low pass filter at 600/300Hz. Events were identified by FFT deconvolution. mEPSC events were excluded based on the following criteria: Amplitude lower than 7 pA. Cells were excluded if the access resistance raised above 20 MOhm before or after the recording. For adult mEPSC we exclude cells if they had a mEPSC frequency above 7 hertz, a membrane resistance below 120 mOhm and a capacitance greater than 60 pF. For P14 mEPSC we just excluded based on access resistance.

For current-clamp analysis. we excluded neurons with a drop in peak AP voltage over the duration of the current step greater than 15 mV, and with and the peak AP voltage lower than 30 mV. Interneurons were excluded based on half-width and firing frequency. Rheobase was defined as the lowest current step that triggerer at least one action potential. The AP threshold was identified using the 3^rd^ derivative. The I-V curve was calculated by fitting a regression between current step amplitude and delta-V for the first 6 current steps. The timing of AP waveform features was the time from spike threshold to the AP feature.

### Statistical analyses

Two-way ANOVA was performed for comparing parameters affected by two factors. All plots show meant +/- SEM. Each shape in a plot group (i.e. Fmr1-/y x D1-SPN) represents a unique mouse for that group. Statistical significance was considered at P < 0.05. Statistical calculations were performed using StatsModels (Python). Specific statistical analyses are detailed in the respective figure legends or results section for each dataset. For the P14 mEPSC experiment a total of 10 D1-SPN and 11 D2-SPN cells from 6 Fmr1+/y mice and 12 D1-SPN and 12 D2-SPN cells from 5 Fmr1-/y mice were analyzed. For the P14 current clamp experiment a total of 20 D1-SPN and 18 D2-SPN cells from 4 Fmr1+/y mice and 11 D1-SPN and 14 D2-SPN cells from 4 Fmr1-/y mice were analyzed. For the adult D1/D2 mEPSC experiment a total of 23 D1-SPN and 28 D2-SPN cells from 4 mice and 30 D1-SPN and 24 D2-SPN cells from 5 mice were analyzed. For the adult D1/D2 current clamp experiment a total of 24 D1-SPN and 16 D2-SPN cells from 5 Fmr1+/y mice and 27 D1-SPN and 14 D2-SPN from 6 Fmr1-/y mice were analyzed. For the adult aripiprazole current clamp experiment a total of 22 cells from 3 Fmr1+/y x Aripiprazole mice, 28 cells from 3 Fmr1+/y x Vehicle mice, 29 cells from 3 Fmr1-/y x Aripiprazole mice, 26 cells from 3 Fmr1-/y x Vehicle mice were analyzed.

---

**Figure S1.**
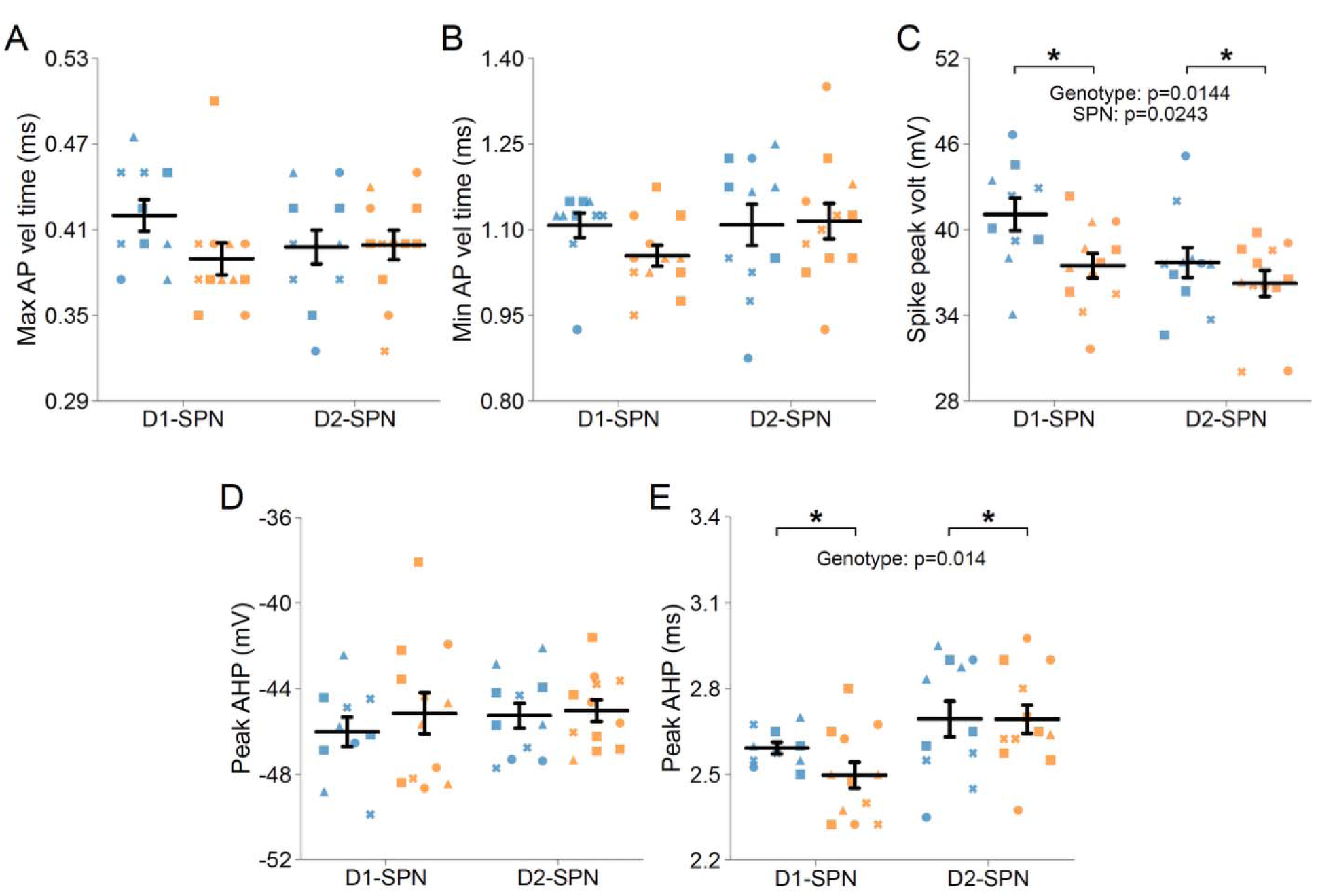
**AP waveform properties in DMS D1- and D2-SPNs of P15 *Fmr1+/y* and *Fmr1-/y mice.*** Fmr1+/y in blue and Fmr1-/y in orange. **(A)** Maximum AP velocity time. **(B)** Time of the inimum AP velocity. **(C)** AP peak voltage. **(D)** Peak AHP voltage. **(E)** Time of the peak AHP voltage. Plots show individual data points, with shape representing a mouse within the group (i.e. Fmr1-/y x D1-SPN) and the summary with mean ± SEM. *P<0.05, **P<0.01, ***P<0.005, ****P<0.001.

**Figure S2.**
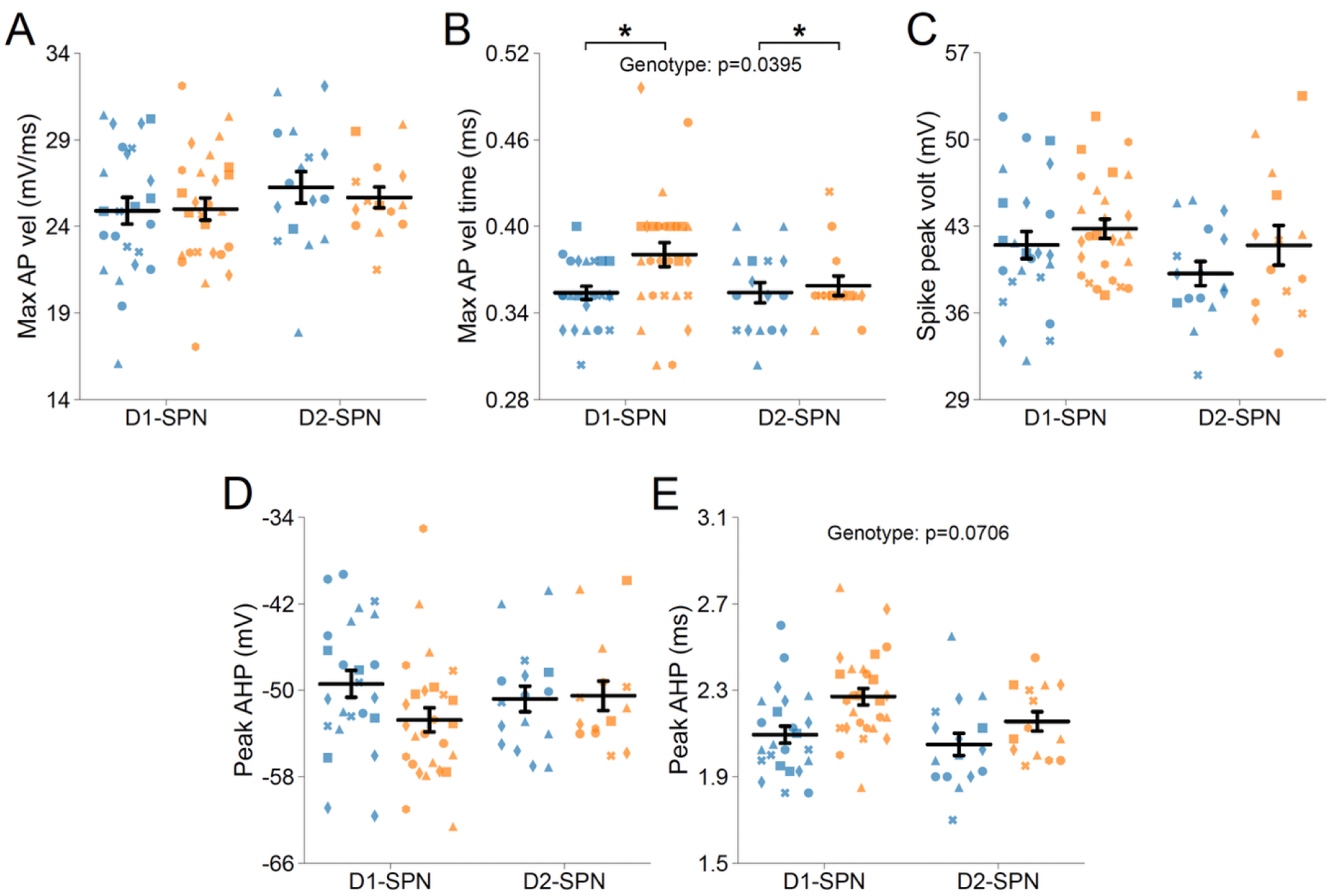
**AP waveform properties in D1- and D2-SPNs of adult P60 *Fmr1+/y* and *Fmr1-/y mice.*** Fmr1+/y in blue and Fmr1-/y in orange. **(A)** Maximum AP velocity. **(B)** Time of the maximum AP velocity. **(C)** AP peak voltage. **(D)** Peak AHP voltage. **(E)** Time of the peak AHP voltage. Plots show individual data points, with shape representing a mouse within the group (i.e. Fmr1-/y x D1-SPN) and the summary with mean ± SEM. *P<0.05, **P<0.01, ***P<0.005, ****P<0.001.

**Figure S3.**
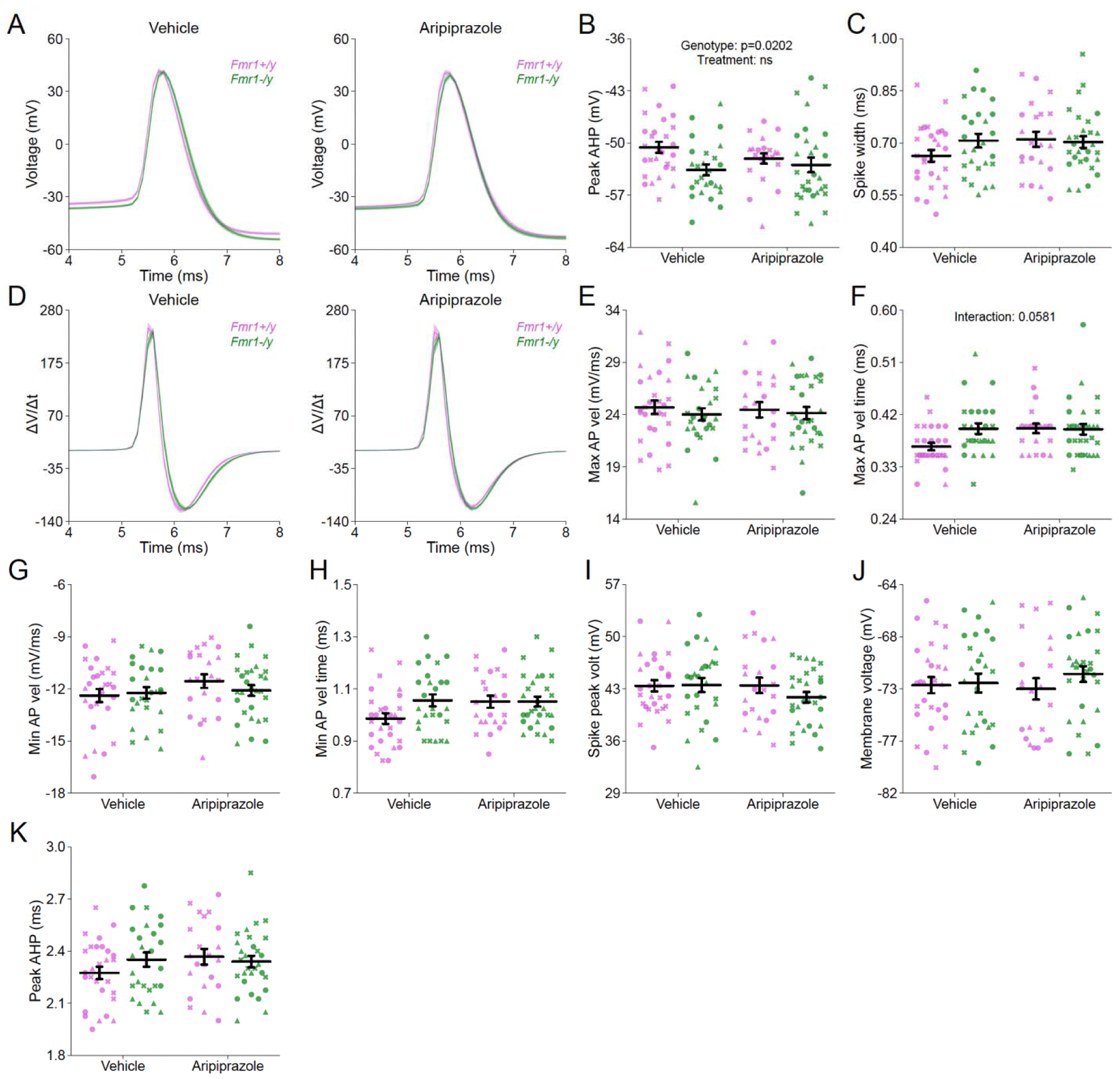
AP waveform properties in vehicle- and aripiprazole-treated adult P60 *Fmr1+/y* and *Fmr1-/y mice.* **(A)** Representative single action potential traces recorded from vehicle- (left) and aripiprazole-treated (right) *Fmr1*+/y (purple) and *Fmr1*-/y (green) mice. **(B)** Peak afterhyperpolarization (AHP) voltage. **(C)** AP half-width. **(D)** First derivative of AP traces (ΔV/Δt) in vehicle- and aripiprazole-treated groups. **(E)** Maximum AP velocity. **(F)** Time of the maximum AP velocity. **(G)** Minimum AP velocity. **(H)** Time of the minimum AP velocity. **(I)** AP peak voltage. **(J)** Resting membrane voltage. **(K)** Second measurement of peak AHP voltage. Plots show individual data points, with shape representing a mouse within the group (i.e. Fmr1-/y x D1-SPN), and summary data presented as mean ± SEM. *P<0.05, **P<0.01, ***P<0.005, ****P<0.001.

